# Age-related alterations in prelimbic cortical neuron *Arc* expression vary by behavioral state and cortical layer

**DOI:** 10.1101/2020.08.11.247262

**Authors:** Abbi R. Hernandez, Leah M. Truckenbrod, Maya E. Barrett, Katelyn N. Lubke, Benjamin J. Clark, Sara N. Burke

## Abstract

Prefrontal cortical and medial temporal lobe connectivity is critical for higher cognitive functions that decline in older adults. Likewise, these cortical areas are among the first to show anatomical, functional, and biochemical alterations in advanced age. The prelimbic subregion of the prefrontal cortex and the perirhinal cortex of the medial temporal lobe are densely reciprocally connected and well-characterized as undergoing age-related neurobiological changes that correlate with behavioral impairment. Despite this fact, it remains to be determined how changes within these brain regions manifest as alterations in their functional connectivity. In our previous work, we observed an increased probability of age-related dysfunction for perirhinal cortical neurons that projected to the prefrontal cortex in old rats compared to neurons that were not identified as projection neurons. The current study was designed to investigate the extent to which aged prelimbic cortical neurons also had altered patterns of *Arc* expression during behavior, and if this was more evident in those cells that had long-range projections to the perirhinal cortex. The expression patterns of the immediate-early gene *Arc* were quantified in behaviorally characterized rats that also received the retrograde tracer cholera toxin B (CTB) in the perirhinal cortex to identify projection neurons to this region. As in our previous work, the current study found that CTB+ cells were more active than those that did not have the tracer. Moreover, there were age-related reductions in prelimbic cortical neuron *Arc* expression that correlated with a reduced ability of aged rats to multitask. Unlike the perirhinal cortex, however, the age-related reduction in *Arc* expression was equally likely in CTB+ and CTB− negative cells. Thus, the selective vulnerability of neurons with long-range projections to dysfunction in old age may be a unique feature of the perirhinal cortex. Together, these observations identify a mechanism involving prelimbic-perirhinal cortical circuit disruption in cognitive aging.

## Introduction

The United States Census Bureau predicts that in less than two decades the number of individuals over the age 65 will exceed the number of children under 18 for the first time in U.S. history (Vespa et al., 2018). Even in the absence of frank neurodegeneration, a majority of people over the age of 65 will suffer from cognitive decline that can impact daily living and the ability to maintain independence (Blazer et al., 2015). Thus, as older adults continue to occupy a larger portion of the population age demographics, understanding the neurobiological underpinnings of cognitive decline becomes increasingly important. It is widely reported that the prefrontal cortex, which is crucial for executive functioning and goal-directed behavior, is particularly susceptible to dysfunction with advancing age (e.g., Barense et al., 2002; Alexander et al., 2008; Morrison and Baxter, 2012; Samson and Barnes, 2013; Banuelos et al., 2014; Beas et al., 2016; Hernandez et al., 2018c). Similarly, structures in the medial temporal lobe, such as the hippocampus (e.g., Rapp and Amaral, 1991; Baxter and Gallagher, 1996; Gallagher et al., 1996; Rosenzweig and Barnes, 2003), and perirhinal cortex (e.g., Liu et al., 2008b; Liu et al., 2008a; Liu et al., 2009; Moyer et al., 2011; Burke et al., 2012; Ryan et al., 2012; Burke et al., 2014; Maurer et al., 2017b; Burke et al., 2018) are known to undergo age-related neurobiological alterations. Importantly, age-related changes in neuron activity patterns and biochemical vulnerabilities are unique between each of these regions. While the cellular changes with age in the prefrontal and perirhinal cortices have been well described, less is known about how aging impacts the functional connectivity between these brain regions. Understanding the interactions between these two regions across the lifespan is critical, as bi-directional communication between the prefrontal and perirhinal cortices supports a number of cognitive functions. For example, disconnecting the perirhinal and prefrontal cortices with pharmacological inactivation or chemogenetic manipulation leads to deficits in biconditional association task performance (Hernandez et al., 2017), sequence memory (Jayachandran et al., 2019), and object-place associations (Barker et al., 2007; Barker and Warburton, 2015).

Our previous work has documented age-related impairments on a biconditional association task in 24 month old male and female rats when compared to rats at 4 months of age (Hernandez et al., 2015; Hernandez et al., 2019a). Functional connectivity between the perirhinal and medial prefrontal cortices is required for biconditional association task performance (BAT; Hernandez et al. 2017), and this circuitry appears to be particularly vulnerable in advanced age (Hernandez et al., 2018b). In fact, when the retrograde tracer cholera toxin subunit B (CTB) was injected into the medial prefrontal cortex to identify neurons in the perirhinal that project to this area, we observed that age differences in the behavior-related expression patterns of the immediate-early gene *Arc* were almost exclusively observed in the neurons that projected to the medial prefrontal cortex. While differences in *Arc* expression were also evident in the prelimbic cortex of the medial prefrontal cortex, whether these occurred in medial temporal lobe-projecting neurons versus other cell types was not determined.

Medial prefrontal cortical neuron activity facilitates task-dependent interactions between the perirhinal and lateral entorhinal cortices (Paz et al., 2007), suggesting that disruptions in prefrontal cortical projections to the perirhinal cortex could impair overall cortical-hippocampal interactions. To determine the extent that aging disrupts activity patterns in the prefrontal cortical neurons that project to the perirhinal cortex, the current study used *Arc* compartmental analysis of temporal activity with fluorescent *in situ* hybridization (catFISH) combined with retrograde labeling. *Arc* is an immediate-early gene that is rapidly transcribed by principal neurons after spiking related to active behavior (Lyford et al., 1995). The transcription kinetics of *Arc* are such that within 1-2 minutes of behavior-related cellular activity, mRNA is evident at transcription loci in the nucleus. Following transcription, 15-20 min after neuron activity, *Arc* is translocated to the cytoplasm (Guzowski et al., 1999, 2001), where the mRNA is locally translated in the dendrites (Steward et al., 1998). Thus, the subcellular location of *Arc* mRNA can be used to infer the activity history of neurons during two distinct behavioral epochs separated by 20 min. Rats also received injections of the retrograde tracer CTB centered on the perirhinal cortex to identify the neurons in the prelimbic cortex that project to this target region. *Arc* expression in all prelimbic neurons and the subset of cells that projected to the perirhinal cortex was then evaluated in association with a cognitive multitask requiring simultaneous working memory and biconditional association retrieval (Hernandez et al., 2019b; Hernandez et al., 2019a) or a control alternation task.

## Methods

### Subjects and Handling

Fisher 344 x Brown Norway F1 (FBN) Hybrid male and female rats from the National Institute on Aging colony at Charles River were used in this study. This rat strain has a longer average life expectancy compared to the inbred Fischer 344 (Turturro et al., 1999), and significant declines in physical function are often not seen until 28 months of age or later (McQuail and Nicolle, 2015). The group that underwent cognitive testing was comprised of young (4-7 mo; male n=8/female n=2) and aged (23-28 mo; male n=5/female n=4) rats.Additional male rats (young n=11, aged n =14) were sacrificed directly from home cages to assess baseline expression of *Arc*, as previously published (Hernandez et al., 2018b).Unfortunately, female rats were not available to include in this control group (https://www.nia.nih.gov/research/dab/aged-rodent-colonies-handbook/oldest-ages-available-nia-aged-rodent-colonies). All rats were housed individually and maintained on a 12-hour reversed light/dark cycle with all behavioral testing occurring in the dark phase.

To encourage appetitive behavior in object discrimination experiments, rats were calorically restricted, receiving 10-30g (1.9 kcal/g) moist chow daily based on rats’ relative body conditions on a scale of 1-5, with 3 being ideal, 1 being emaciated, and 5 being morbidly obese (Hickman and Swan, 2010). Water was provided *ad libitum*. Rats’ body weights were recorded daily, and body condition was assessed and recorded weekly throughout the period of restricted feeding. The body condition score was assigned based on the presence of palpable fat deposits over the lumbar vertebrae and pelvic bones (Hickman and Swan, 2010). Animals with a score under 2.5 were given additional food to promote weight gain. All procedures were performed in accordance with the National Institutes of Health Guide for the Care and Use of Laboratory Animals and approved by the Institutional Animal Care and Use Committee at the University of Florida.

### Behavioral Tasks

Rats began training on a Figure-8 maze for continuous alternation testing. Rats were trained to traverse the maze, making alternating left and right turns after walking the length of the center arm, receiving a Froot Loop reward when correct. After alternating at or above 85% correct for two consecutive days, rats began behavioral testing on the working memory biconditional association task (WM/BAT). WM/BAT necessitates the use of location information (i.e., right or left arm) to select the correct object between a pairwise discrimination. The rat was required to push the object to access a Froot Loop reward in the well beneath. For the first 10 trials on the first day of testing, wells are only partially covered by the objects to encourage learning. When an incorrect object was chosen or the animal made an incorrection alternation (i.e. two turns in the same direction), the objects and reward were removed, and the animal proceeded to the next trial.

### CTB Retrograde Tracer Injection Surgeries

After behavioral training, rats underwent stereotactic surgery to inject the retrograde tracer cholera toxin subunit B (CTB) Alexa Fluor 594 Conjugate (CTB594; catalog #C22842) into the right perirhinal cortex (PER). Under isoflurane anesthesia (1%–3%), an incision was made to expose Bregma. A small craniotomy was made at −5.5 mm for young animals and −5.7 mm for aged animals posterior to Bregma and +7.2 mm for young animals and +7.4 mm for aged animals lateral to Bregma. Coordinates for old animals were slightly adjusted based on differences in the length between bregma and lambda between age groups (see Figures 1A-C for representative injections). Using a glass pipette backfilled with CTB594, a Nanoject II Auto-Nanoliter Injector (Drummond Scientific Company) was used to inject 100 nL of CTB via two 50 nL infusions at −4.5 mm for young animals and −4.64 mm for aged animals ventral to the brain surface. After waiting 60 seconds, the pipette was moved up 100 μm and another two 50 nL volumes of CTB were injected, and this process was repeated for a total of six injections. Therefore, the final injection was given at a depth of −4.3 mm and −4.44 mm ventral to the brain surface, respectively. After the final injection, the pipette was left in place for 150 seconds and then slowly advanced up by 0.5 mm, where it was left in place for another 150 seconds allowing for the tracer to diffuse away. After the final waiting period, the pipette was slowly removed from the brain. During surgery and postoperatively, the nonsteroidal anti-inflammatory meloxicam (Boehringer Ingelheim Vetmedica, Inc., St. Joseph, MO; 1.0 mg/kg subcutaneously) was administered as an analgesic daily for 2 days. All animals were given 7 days to recover before resuming behavioral testing.

**Figure 1:**
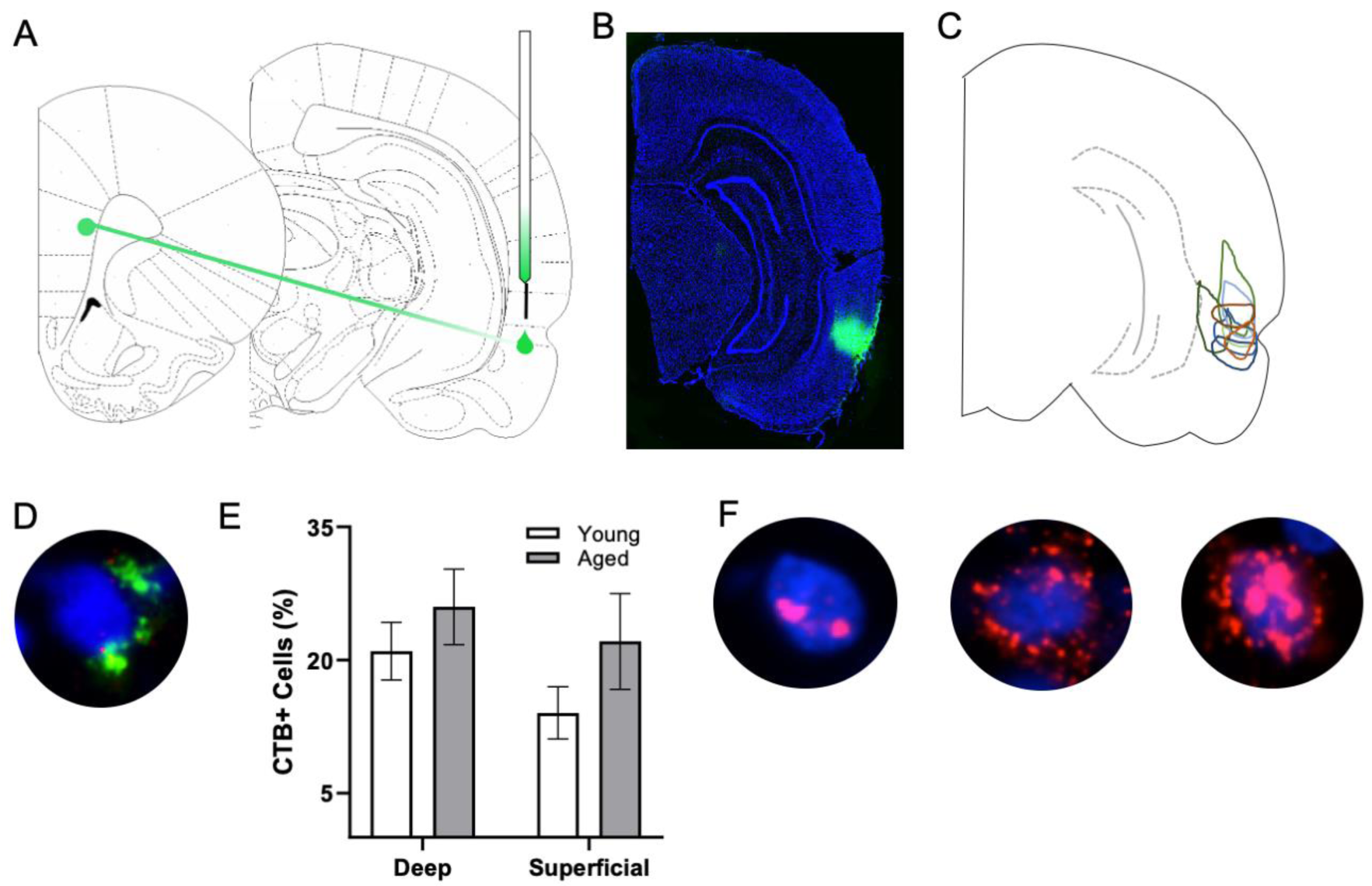
*Arc* and CTB labeling. **(A)** Schematic of target site for CTB injection surgery. **(B)** Representative image of CTB spread. **(C)** Diagram of CTB injection spread for each subject. **(D)** Example image of CTB positive cell. **(E)** Percentage of CTB positive cells in deep and superficial prelimbic cortex did not significantly differ between young and aged rats in either the deep (t_[17]_ = 0.95; p = 0.36) or superficial layers (t_[16]_ = 1.38; p = 0.19).**(F)** Example images of *Arc* positive cells (from left to right) with *Arc* foci only, cytoplasm only, and both foci and cytoplasm.

### Sacrifice and Tissue Collection

On the day of sacrifice, a timed behavioral experiment was performed for *Arc* cellular compartment analysis of temporal activity with fluorescent *in situ* hybridization (catFISH) (Guzowski et al., 1999). Thirty minutes prior to sacrifice, rats performed either the WM/BAT or alternation task for 5 minutes each in 2 distinct behavioral epochs separated by a 20-minute rest in their home cage. The order of behavioral testing was counterbalanced across rats. This design enabled neurons activated during the two different behaviors to be differentiated based on the subcellular location of *Arc* mRNA (Guzowski et al., 1999).

Following the second epoch of behavior, rats were immediately placed into a bell jar containing isoflurane-saturated cotton (Abbott Laboratories, Chicago, IL, USA) separated from the animal by a wire mesh shield. Animals rapidly lost the righting reflex (<30 seconds), after which they were immediately euthanized via rapid decapitation. Tissue was extracted and flash frozen in 2-methylbutane (Acros Organics, NJ, USA) chilled in a bath of dry ice with 100% ethanol (∼−70°C). Tissue was stored at −80°C until cryosectioning. Tissue blocks were made that contained brains from each experimental group to control for slight variations in tissue labeling across slides. These tissue blocks where sectioned at 20 μm on a cryostat (Microm HM550) and thaw mounted on Superfrost Plus slides (Fisher Scientific). Sliced tissue was stored at −80°C until *in situ* hybridization.

### In Situ Hybridization

*In situ* hybridization was performed to label *Arc* mRNA for catFISH as previously described (Guzowski et al., 1999). Briefly, a commercial transcription kit and RNA labeling mix (Ambion REF #: 11277073910, Lot #s: 26591021, 26591020; Austin, TX) were used to generate a digoxigenin-labeled riboprobe using a plasmid template containing a 3.0 kb *Arc* cDNA. After tissue was incubated with the probe overnight, *Arc*-positive cells were detected with anti– digoxigenin-HRP conjugate (Roche Applied Science Ref #: 11207733910, Lot #: 10520200; Penzberg, Germany). Fluorescein (Fluorescein Direct FISH; PerkinElmer Life Sciences, Waltham, MA) was used to visualize labeled cells, and nuclei were counterstained with DAPI (Thermo Scientific).

### Visualization and analysis

After staining, z-stacks at 1 µm increments were collected by fluorescence microscopy on a Keyence BZX-810 digital microscope (Keyence Corporation of America, Itasca, IL). Three images were taken from the deep and superficial prelimbic cortex. For one aged rat, only 2 images from superficial prelimbic cortex were usable due to tissue damage. Experimenters blind to age and order of behavioral tasks quantified *Arc* expression in neurons using ImageJ software with a custom written plugin for identifying and classifying cells (available upon request). Only cells that were completely within the frame of the image and within the median 20% of the optical planes were included. The total number of cells were identified before the CTB and *Arc* color channels were visible so that signal in these channels did not bias initial segmentation results. Cells were classified as positive for nuclear *Arc*, cytoplasmic *Arc*, both nuclear and cytoplasmic *Arc*, or negative for *Arc* as well as positive or negative for CTB. A cell was considered *Arc* nuclear positive if 1 or 2 fluorescently labeled foci were visible within the nucleus on at least 4 consecutive planes of the z-stack. A cell was counted as *Arc* cytoplasmic positive if fluorescent labeling could be detected above background surrounding at least 1/3 of the nucleus on 2 adjacent planes. Cells meeting both these criteria were counted as both *Arc* nuclear and cytoplasmic positive. *Arc* in the cytoplasm indicates that the cell fired during epoch 1, as it takes approximately 20-30 minutes for the mRNA to translocate to the cytoplasm. *Arc* foci in the nucleus indicate that the cell fired during epoch 2. Figure 1F shows representative examples of the subcellular distribution of *Arc*. Cells were considered to be positive for CTB if at least 1/3 of the nucleus was surrounded by labeling on at least 2 adjacent planes (see figure 1D for example). There were no differences in the proportion of CTB labeled cells across young and aged rats for either the deep (t_[17]_ = 0.95; p = 0.36) or superficial layers (t_[16]_ = 1.38; p = 0.19;

### Statistical Analysis

Differences in neuronal activation during the behavioral epochs in relation to age, cortical layer and CTB status were statistically evaluated by calculating the mean percentage of *Arc* positive cells per rat as done previously (e.g., Hartzell et al., 2013; Hernandez et al., 2018b). This avoids inflating statistical results by including multiple measures from one animal and does not violate the assumption of independent observations (Aarts et al., 2014). All behavior was counterbalanced across rats to avoid bias from cytoplasmic versus nuclear foci labeling. Potential effects of age, cellular layer and task were examined using repeated measures ANOVAs (ANOVA-RM) with the within-subject factors of behavioral task, cortical layer or CTB status, and the between-subjects factor of age. All analyses were performed with the Statistical Package for the Social Sciences v25 (IBM, Armonk, NY) or GraphPad Prism version 7.03 for Windows (GraphPad Software, La Jolla, California USA). Statistical significance was considered at p-values less than 0.05.

## Results

### Baseline Arc Expression in Caged Control Rats

Baseline *Arc* expression was quantified in the superficial and deep layers of the prelimbic region of the medial prefrontal cortex from a subset of animals sacrificed directly from their home cage that were not behaviorally trained. Baseline *Arc* expression was significantly higher in the deep compared to superficial layers (F_[1,23]_ = 4.80; p = 0.04). There were no differences, however, across age (F_[1,23]_ = 0.58; p = 0.46). Moreover, age and layer did not significantly interact (F_[1,23]_ = 0.82; p = 0.38). Post hoc analyses for the deep and superficial layers individually, indicated that while there was no age difference within the deep layers (t_[23]_ = 0.03; p = 0.98; Figure 2A), there was a trend for aged rats to have higher *Arc* expression within the superficial layers (t_[23]_ = 1.977; p = 0.06; Figure 2B). This observation is consistent with a previous study that reported a higher proportion of cells with *Arc* in the superficial layers of prelimbic cortex in aged compared to young rats (Hernandez et al., 2018b).

**Figure 2:**
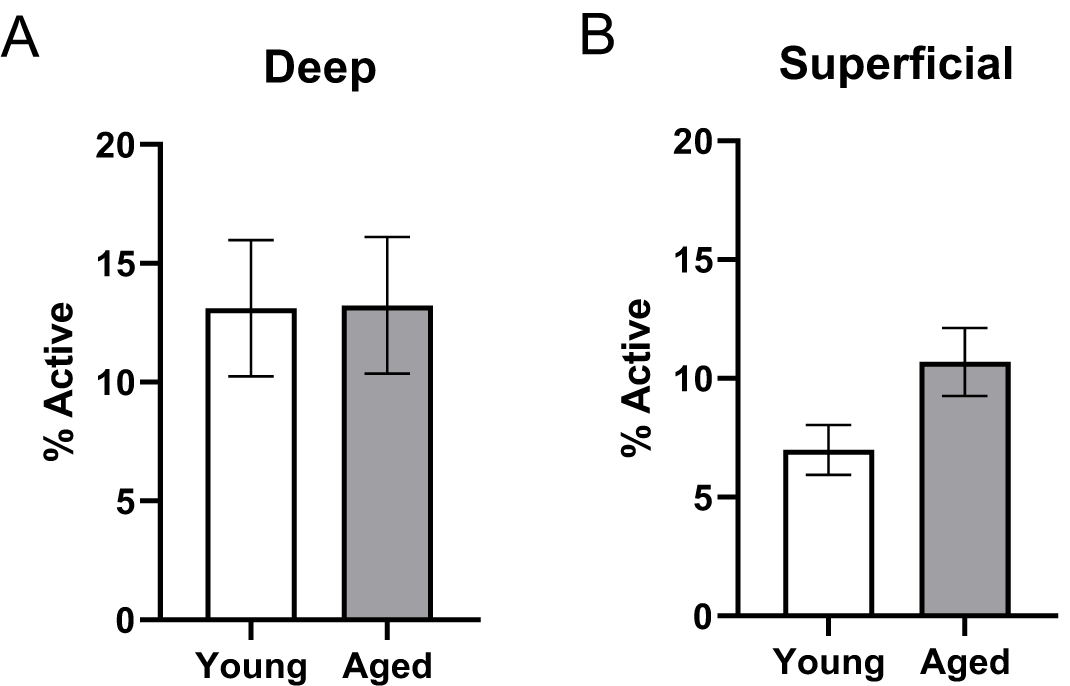
Quantification of *Arc* expression at baseline. **(A)** There were no differences in the percentage of active cells in the deep layers of the prelimbic cortex (t_[23]_ = 0.03; p = 0.98; Figure 2A) in animals sacrificed directly from the home cages. **(B)** In the superficial prelimbic cortical layers, there was a trend for aged rats to have higher baseline *Arc* expression within the superficial layers (t_[23]_ = 1.977; p = 0.06). All data represent group means ± 1 SEM.

### Aged rats performed worse than young rats on WM/BAT testing

It has been widely reported that aged rats are impaired at acquiring a biconditional association task rule compared to young animals (Hernandez et al., 2015; Hernandez et al., 2018b; Hernandez et al., 2019b; Hernandez et al., 2019a). In line with these previous data, on the day of the experiment aged rats made significantly more object selection errors than the young rats (t_[18]_ = 3.11; p = 0.006; Figure 3A), despite comparable days of training. Conversely, on the final day of testing there was not a significant age difference in ability to correctly alternate (t_[18]_ = 0.24; p = 0.81). Importantly, the number of trials completed during the two behavioral epochs on the day that tissue was collected for catFISH did not significantly differ between young and aged rats for either WM/BAT or the alternation task (p > 0.15 for both tasks; Figure 3B).

**Figure 3:**
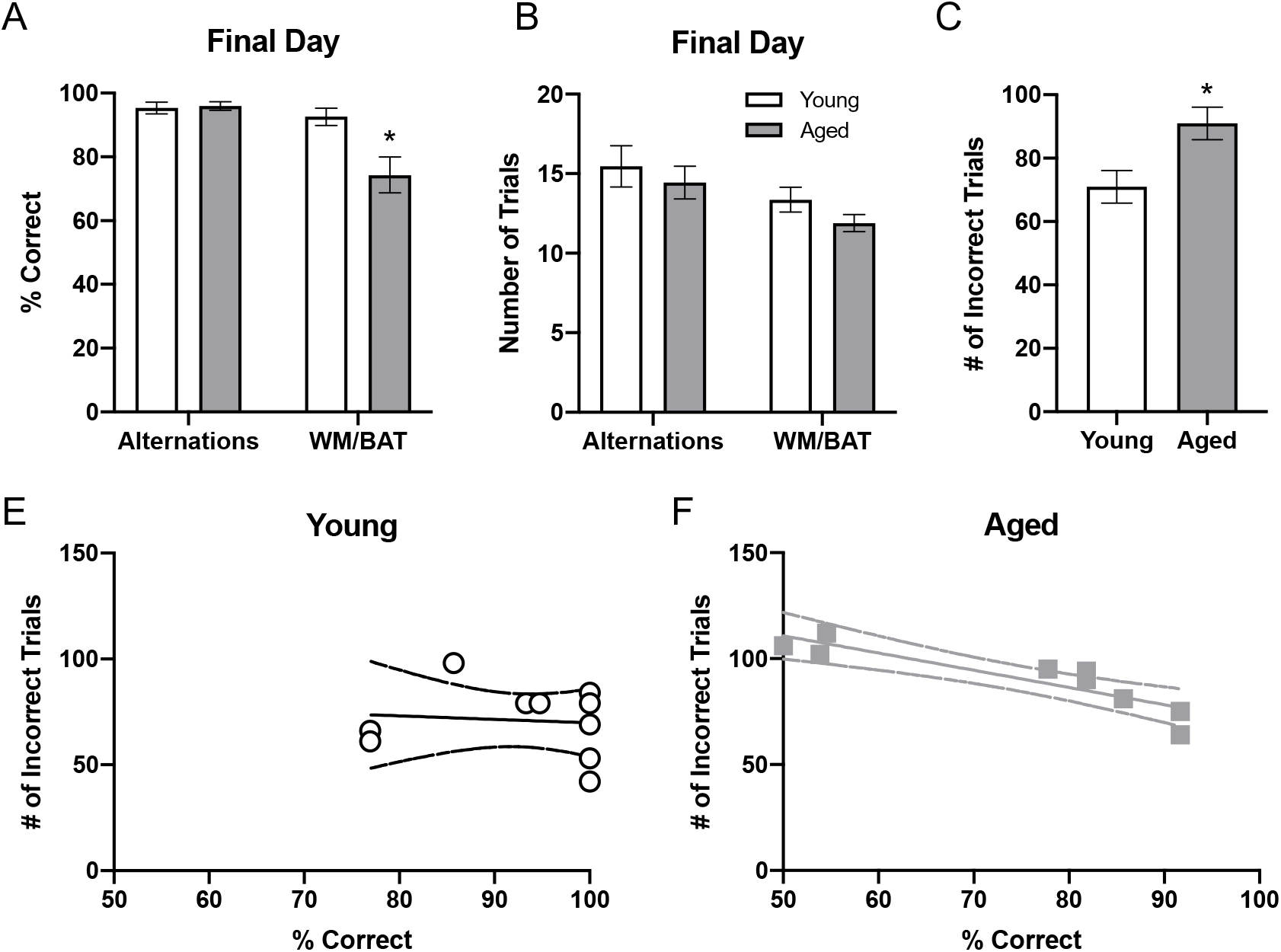
Behavioral performance in young and aged rats. **(A)** On the final day of testing (the day of sacrifice), the aged rats’ performances was significantly worse than that of the young animals (t_[18]_ = 3.11; p = 0.006). In contrast, there was not a significant age difference in alternation performance (t_[18]_ = 0.24; p = 0.81). **(B)** There were no significant differences in the number of trials completed on the day of sacrifice on either alternations or WM/BAT (p > 0.15 for both tasks). **(C)** The total number of incorrect trials during WM/BAT training was significantly greater in the aged compared to young rats (t_[18]_ = 2.71, p < 0.02). **(D)** In young rats, the number of incorrect trials did not correlate with WM/BAT performance on the final day of testing (R_[8]_ = −0.02; p = 0.96). **(E)** In aged rats, there was a significant correlation between errors during training and performance on the last day (R_[7]_ = −0.90, p < 0.01).

To relate performance on the day that tissue was collected to overall WM/BAT acquisition abilities, the total number of incorrect trials during training was tabulated and compared with their performance on the final day of testing. Aged rats made significantly more errors than the young animals during training (t_[18]_ = 2.71, p < 0.02; Figure 3C). For the young rats (Figure 3D), there was no significant correlation between the training data and final performance (R_[8]_ = −0.02; p = 0.96), likely due to near ceiling level performance on the final day of testing. In contrast, for the aged rats there was a significant correlation between the number of incorrect trials during all training and performance on the last day (R_[7]_ = −0.90, p < 0.01). These data indicate that WM/BAT performance on the day of tissue collection was likely representative of good and bad performers within the aged group across all behavioral training.

### Behaviorally induced Arc Expression

Figures 4A/B show the average population activity within superficial and deep layers of the prelimbic cortex during both alternation and WM/BAT epochs. ANOVA-RM on the non-normalized percentage of *Arc* positive cells comparing the 2 tasks with the between subjects factors of age and region did not show a main effect of task (F_[1,34]_ = 0.16; p = 0.69), similar to previous data (Hernandez et al., 2018b). Across both tasks and layers, the proportion of active cells during behavior was significantly reduced in the aged rats (F_[1,34]_ = 4.29; p < 0.05). Moreover, the proportion of active cells was higher in the deep compared to superficial layers (F_[1,34]_ = 6.56; p < 0.02). Age group and prelimbic cortical layer, however, did not significantly interact (F_[1,34]_ = 0.10; p = 0.77). Furthermore, task did not significantly interact with either age or layer (p > 0.60 for both comparisons).

**Figure 4:**
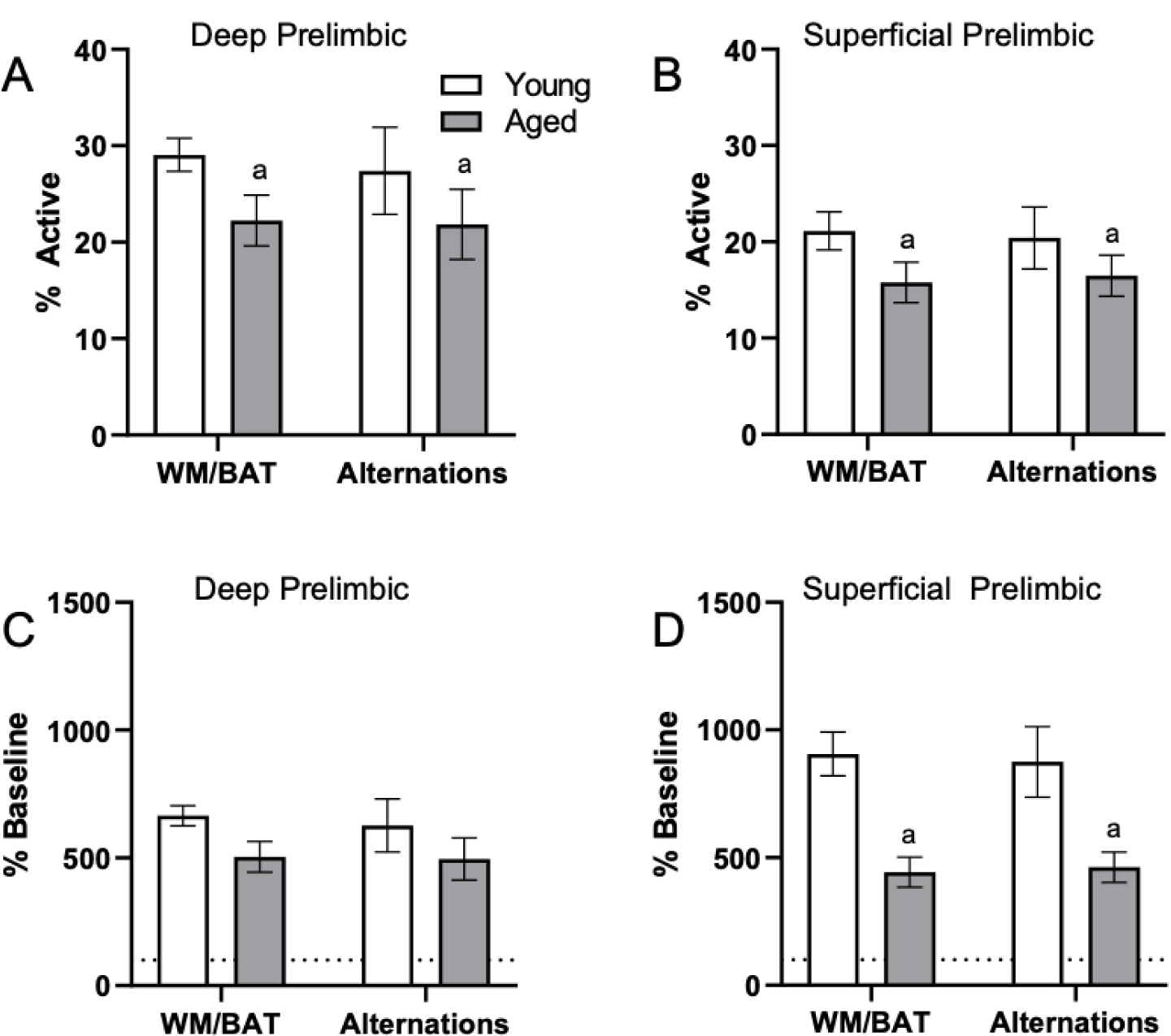
Prelimbic cortical *Arc* expression during behavior. Behaviorally induced *Arc* expression shown as percentage of active cells within the (A) deep or (B) superficial layers of the prelimbic cortex during alternation and WM//BAT epochs (X axis) was significantly reduced in aged compared to young rats (F_[1,34]_ = 4.29; p < 0.05). Percent increase in *Arc* expression from baseline in the (C) deep and (D) superficial layers normalized to home cage control data. There was a significant effect of age in the superficial layers (F_[1,34]_ = 14.54; p < 0.001). All data represent group means ± 1 SEM; ^a^ indicates a significant main effect of age.

Because aged rats show a trend towards elevated proportions of cells that were positive for *Arc* expression within the home cage in superficial prelimbic cortex, *Arc* expression was normalized relative to rats sacrificed directly from the home cages to quantify the extent that behavior elevated prelimbic activity patterns above baseline conditions (Fletcher et al., 2014; Pereira et al., 2015). ANOVA-RM on the normalized percentage of *Arc* positive cells with the within subjects factor of task (WM/BAT versus alternation) and the between subjects factors of age and cortical layer did not detect a significant main effect of task on the increase from baseline expression (F_[1,34]_ = 0.15; p = 0.70). The aged rats, however, had a significantly lower increase from baseline compared to the young animals (F_[1,34]_ = 14.54; p < 0.001). In contrast to the raw data, there was not a significant difference in the percent increase from baseline between the deep and superficial layers (F_[1,34]_ = 1.66; p = 0.21). This suggests that the higher levels of activity in the deep compared to superficial layers of the prelimbic cortex were due to differences in activity at baseline, but the neurons in the different layers were comparably engaged by behavior. When the deep and superficial layers were examined individually, aged rats trended towards significantly fewer active cells than young within the deep layers (F_[1,34]_ = 3.73; p = 0.06; Figure 4C), and this reached significance within the superficial layers (F_[1,34]_ = 21.47; p < 0.01; Figure 4D) of the prelimbic cortex. Task did not significantly influence activity within either region (p > 0.75 for both regions), nor did task significantly interact with age (p > 0.79 for both regions).

### Neuronal Arc expression correlates with behavioral performance in aged rats

Behavioral performance on the final day of testing was compared to the proportion of active cells within deep and superficial layers of the prelimbic cortex for the alternation and WM/BAT tasks separately. There was no correlation between *Arc* expression and alternation task performance in either the young or aged rats (p > 0.21 for all comparisons; Figure 5A-B). The lack of correlation in either age group or cell layer was likely due to the animals alternating with a high degree of accuracy by this time point in the experiment. Interestingly, a different pattern was observed when comparing the percent of cells positive for *Arc* to behavioral performance during the WM/BAT. Within the deep layers of the prelimbic cortex, there was a significant positive correlation between performance and the percentage of *Arc* positive cells in the aged rats (R_[7]_ = 0.73; p < 0.04; Figure 5C). This was not observed in the young animals or in the superficial layers for either age group (p > 0.56 for all comparisons; Figure 5D).

**Figure 5:**
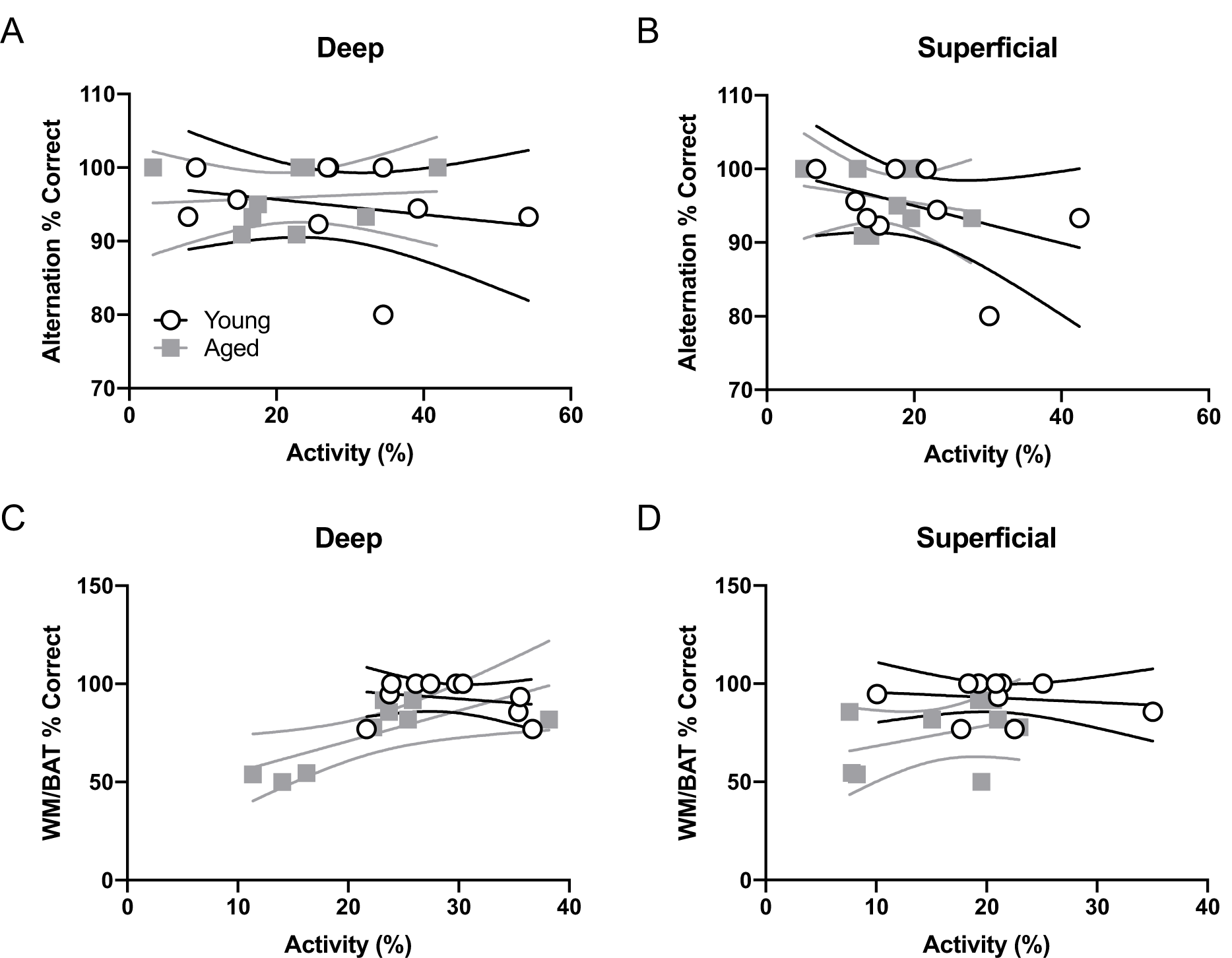
Percent of active cells and behavioral performance. There was no correlation between the percent of neurons expressing *Arc* and alternation task performance in young or aged rats for the **(A)** deep or **(B)** superficial layers of the prelimbic cortex (p > 0.21 for all comparisons). **(C)** In the aged (R_[7]_ = 0.73; p = 0.04), but not young group (R_[8]_ = −0.24; p = 0.53), rats with more *Arc*-positive cells had significantly better WM/BAT performance. **(D)** The percent of *Arc*-positive cells did not correlate with WM/BAT performance in either age group in the superficial layers (p > 0.56 for both groups).

### Arc expression within prelimbic cortical neurons that project to the perirhinal cortex

To investigate age-related changes in neuronal activity in the subset of prelimbic neurons that project to the perirhinal cortex (PER), analysis of behaviorally induced *Arc* expression within the prelimbic cortex was restricted to neurons that were CTB+. As the CTB was injected into the PER, neurons in the prelimbic cortex that are CTB+ are the subset of cells that project directly to this region. As previously published for the reciprocal projection from PER to prelimbic cortex, the proportion of *Arc*+ cells appeared to be higher in the CTB+ cells compared to the whole population. Thus, the proportion of CTB-cells expressing *Arc* relative to CTB+ cells were compared using ANOVA-RM across tasks and age groups. There were significantly more cells expressing *Arc* in the CTB+ compared to the CTB-neuron populations across both tasks (F_[3,102]_ = 27.06; p < 0.01; Figure 6A-B), and there was a significantly greater number of active CTB+ cells within the deep relative to superficial layers of the prelimbic cortex (F_[1,34]_ = 5.05; p < 0.03). Furthermore, cortical layer significantly interacted with CTB status and *Arc* expression (F_[3,102]_ = 3.55; p < 0.02) because the difference in the percent of *Arc*+ neurons between CTB+ and CTB-cells was greater in the deep compared to superficial layers. Age did not influence CTB/*Arc* expression patterns across the two tasks (F_[1,34]_ = 2.27; p = 0.14), nor did age significantly interact with any other variables (p > 0.56 for all comparisons).

**Figure 6:**
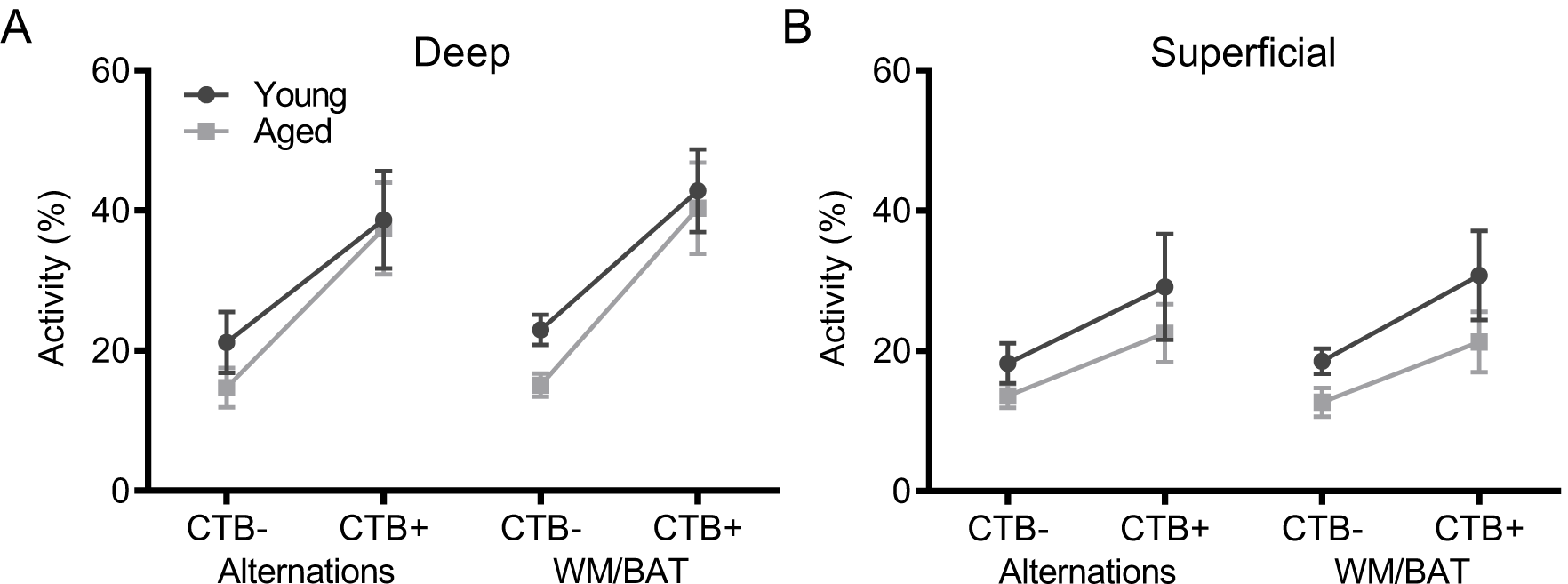
*Arc* expression in prelimbic cortical neurons that project to the perirhinal cortex. For both **(A)** deep and **(B)** superficial layers, significantly more cells expressed *Arc* in the CTB+ cells compared to the CTB-cells (F_[3,102]_ = 27.06; p < 0.01), although this difference was greater in deep layers. All data represent group means ± 1 SEM.

To investigate whether cellular activity within PER projecting neurons of the prelimbic cortex correlated with behavioral performance, the percentage of CTB+ neurons that were also *Arc*+ were plotted against behavioral performance. In the aged rats, WM/BAT performance was significantly correlated with the percentage of CTB+ cells that expressed *Arc* during behavior in the deep prelimbic cortical layers (R_[7]_ = 0.73, p < 0.02) and trended toward a significant correlation in the superficial layers (R_[7]_ = 0.65, p < 0.06). The relationship between WM/BAT performance the percent of CTB+ cells that expressed *Arc* did not reach statistical significance in the young group for either the deep or superficial layers (p > 0.73 for both comparisons).

### Similarity scores do not differ by age within the prelimbic cortex

To determine the degree to which population activity overlap for the two different behavioral tasks was influenced by age group and cortical layers, a similarity score was calculated (Vazdarjanova and Guzowski, 2004). The similarity scores between behavioral epochs 1 and 2 were calculated as:

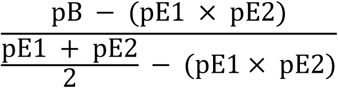

where pB is the proportion of cells active during both epochs and pE1 and pE2 are the proportion of cells active during epochs 1 and 2 respectively. Figure 8 shows the similarity scores for each region. There were no significant age differences in either deep (t_[17]_ = 0.52; p = 0.62) or superficial layers (t_[17]_ = 0.60; p = 0.56; Figure 8) of prelimbic cortex. This observation is different from our previously published data, in which the aged rats had a significantly lower similarity score compared to young animals across an object-place association and easier alternation task. This apparent discrepancy could be due to the enhanced working memory component in both WM/BAT and alternation task used here compared to the tasks used in Hernandez et al., (2018b). Additionally, it is also conceivable that differences in the amount of training between the two experiments contributed to differences in similarity score outcomes across age groups. Specifically, in Hernandez et al. (2018b) all rats were trained to a criterion performance, which was not the case in the present study. These data suggest that there was a degree of population overlap across tasks in both age groups, although a subset of cells were task selective.

**Figure 7:**
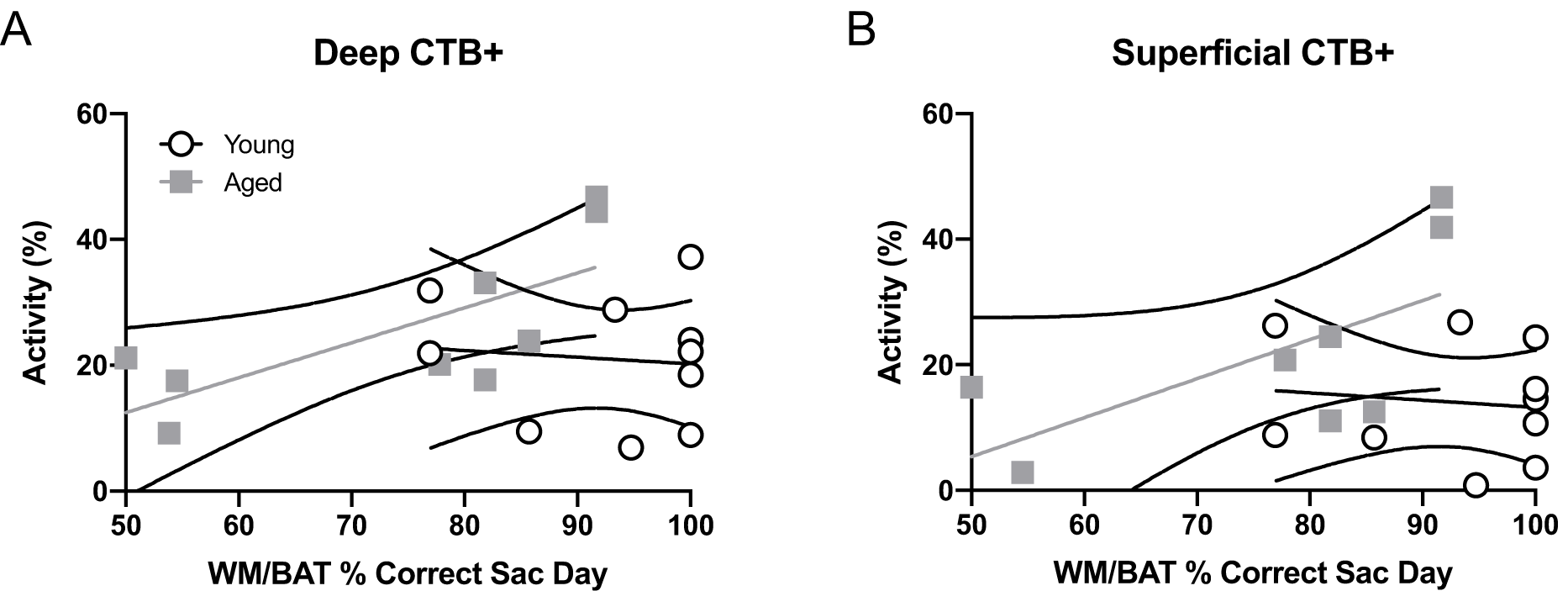
Projection neuron *Arc* expression and WM/BAT performance. When cellular activity was examined in the CTB+ cells, aged rats had a significant correlation between activation and WM/BAT performance in the **(A)** deep layers of prelimbic cortex (R_[7]_ = 0.73, p < 0.02). **(B)** In the superficial layers of prelimbic cortex the relationship between activation and WM/BAT performance trended towards significance (R_[7]_ = 0.65, p < 0.06). In the young rats, however, prelimbic activation did not correlate with behavioral performance in either the **(A)** deep or **(B)** superficial layers. (p > 0.73 for both comparisons)

**Figure 8:**
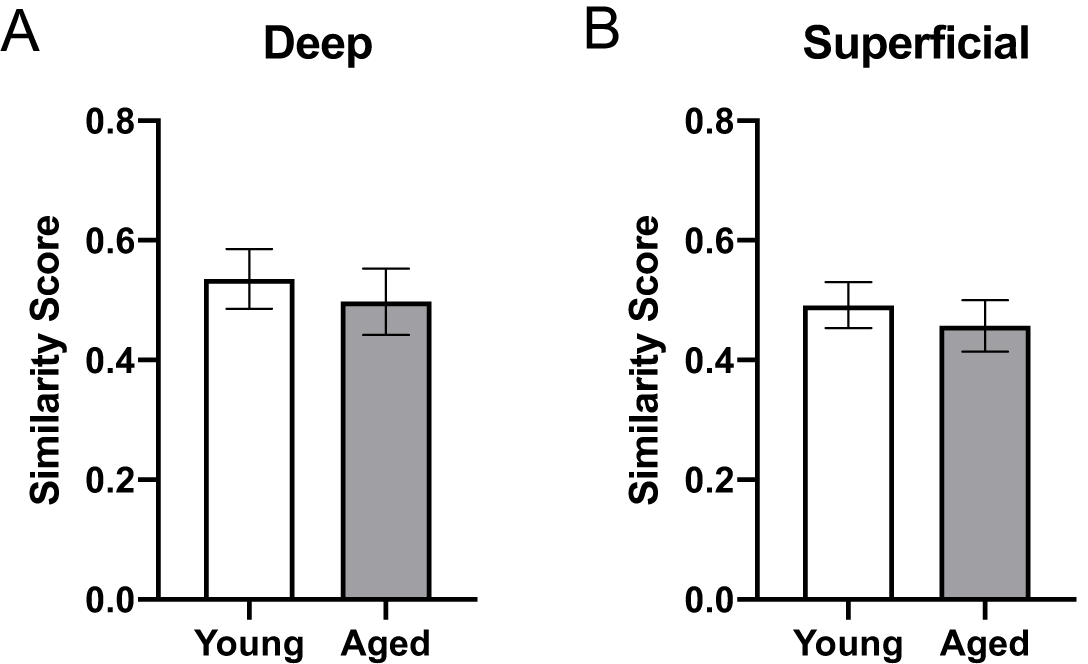
Ensemble activity overlap across behavioral tasks. Similarity scores across the two behavioral epochs did not differ across young and aged rats for both the **(A)** deep and **(B)** superficial layers of the prelimbic cortex, indicating different networks were activated across the two tasks in both groups (p > 0.5 for all comparisons). All data represent group means ± 1 SEM.

## Discussion

The current study examined behavior-related principal neuron activation in anatomically defined circuits by using the cellular compartment analysis of *Arc* expression combined with retrograde labeling to identify prelimbic cortical neurons that projected to the perirhinal cortex.

Here we report several observations supporting the idea that bi-directional functional connectivity between the prelimbic and perirhinal cortices is critical for higher cognitive function and vulnerable in advanced age. Replicating previous work (Hernandez et al., 2018b), aged rats in the current study had a higher percentage of neurons in the superficial layers of prelimbic cortex that expressed *Arc* when rats were in the home cages compared to young animals (Fig 2B). Moreover, also consistent with previous work (Hernandez et al., 2015; Hernandez et al., 2018a; Hernandez et al., 2019a), aged rats were impaired at performing a working memory/biconditional association task (WM/BAT; Fig 3), which requires functional connectivity between the prelimbic and perirhinal cortices (Hernandez et al., 2017).

During behavior on both the WM/BAT and continuous alternation task, the aged rats had reduced expression of *Arc* in both the deep and superficial layers of the prelimbic cortex compared to young animals. A previous publication reported that when aged rats were trained on a biconditional association task (similar to the task used here) until they reached the same performance criteria as young rats, there was increased activation of neurons in the deep layers of the prelimbic cortex and decreased activation in the superficial layers in aged compared to young rats (Hernandez et al., 2018b). In the current study, rats received fewer days of training on the WM/BAT. Thus, when *Arc* expression patterns were evaluated, there was a significant performance deficit in the aged rats in contrast to the matched performance across age groups in Hernandez et al. (2018b). Under these behavioral circumstances, aged rats had lower activation than young animals in both the superficial and deep layers of the prelimbic cortex when the total percent of *Arc*+ cells was evaluated and when the percent increase from baseline was quantified (Fig 4). This finding has two important implications. First, it appears that aged behaviorally impaired rats are less able to recruit neurons in the prelimbic cortex in support of cognitive function. Second, it suggests that the recruitment of neurons in the deep layers of prelimbic cortex is a critical mechanism for supporting WM/BAT performance. This hypothesis is bolstered by the novel observation that aged rats with better WM/BAT performance had a higher percentage of cells that expressed *Arc* in the deep layers of the prelimbic cortex (Fig 5). Notably, the aged rats with the best performance had activation levels in the prelimbic cortex that were comparable to the young group. This observation suggests that maintenance of prelimbic cortical function in advanced age, rather than compensation, was associated with better behavioral output (Barulli and Stern, 2013).

Notably, age-associated differences in the proportion of neurons that expressed *Arc* were different between quiet and active behavioral states. In the home cage, that is, under baseline conditions, aged rats had more neurons that were positive for *Arc* mRNA in the superficial layers of prelimbic cortex. In contrast, aged rats had fewer neurons that were active during behavior for both the WM/BAT and continuous alternation tasks. This is consistent with a previous study that showed reduced levels of *Arc* protein in the prelimbic cortex of aged compared to young rats following behavior on a T-maze (Pereira et al., 2015). Elevated activation within the prefrontal cortex in the absence of a cognitive load that does not appropriately elevate to match increasing behavioral demands is consistent with the idea that the aged prefrontal cortex and its associated cortical networks have a reduced dynamic range of neural activation (Kennedy et al., 2017; Rieck et al., 2017). Mechanistically, reductions in the dynamic range of prefrontal cortical activity could result from a disrupted balance between inhibition and excitation that has been reported for this region (Bories et al., 2013; Banuelos et al., 2014; McQuail et al., 2015; Carpenter et al., 2016).

The observation that aged rats were less able to recruit additional resources as cognitive load increased is also consistent with the Compensation-Related Utilization of Neural Circuits hypothesis (CRUNCH). CRUNCH postulates that more neural resources are recruited by older adults (or animals) during tasks with minimal cognitive load; in this case, sitting in the home cage. This increased activation, could serve to compensate for a network that is compromised. As tasks become more difficult, the limits of network capacity may be reached in older adults and these compensatory mechanisms are no longer effective, leading to less activation relative to younger subjects (Reuter-Lorenz and Cappell, 2008; Grady, 2012). Older subjects that are able to maintain patterns of network activation across different behavioral states that resemble that of their younger counterparts may, therefore, might be better able to sustain cognitive performance.

The prelimbic cortex sends dense projections to both PER subdivisions areas 35 and 36, which originate from both deep and superficial cortical layers (Burwell and Amaral, 1998; Hwang et al., 2018). Dense retrograde labeling across cortical layers was also observed in the current experiment, and this did not vary between the young and old age groups. Reciprocal fibers connecting the prelimbic and perirhinal cortices travel through the parahippocampal branch of the cingulum (Bubb et al., 2018). While the integrity of the cingulum across the lifespan has not been assessed in rats, diffusion tensor imaging data from monkeys (Makris et al., 2007) and humans (Bennett et al., 2015; Bennett and Stark, 2016) suggests that some branches of the cingulum undergo age-related declines in tract integrity. The observation that CTB labeling did not vary across age groups, suggests that age-associated declines in cingulum integrity are due to disrupted myelin rather than a loss of fibers. It is also worth noting that the structural and functional connectivity between brain networks that send axons through the cingulum (Hirsiger et al., 2016) have been reported to not correlate in older adults. This finding further corroborates the notion that potential age-related declines in cingulum integrity are not likely to account for the age differences observed here.

When *Arc* expression was evaluated in prelimbic to perirhinal cortical projection neurons, this cell population was more likely to express *Arc* than those cells that were not labeled with the retrograde tracer cholera toxin B (CTB). The increased probably of activation among projection neurons is consistent with data from projection neurons in CA1 of the hippocampus (Hernandez et al., 2018b), lateral entorhinal (Maurer et al., 2017a), and perirhinal cortices (Hernandez et al., 2018b). Interestingly, the activation difference between CTB positive and negative cells was more pronounced in the deep compared to superficial layers. This is perhaps not surprising giving the anatomical differences between the layers and overall higher levels of activation in the deep layers. A higher probability of activation in projection neurons may be related to the physiological observation that a subset of cells are more likely to fire across different experiences (Mizuseki and Buzsáki, 2013; Buzsáki and Mizuseki, 2014) and a small world network organization (Watts and Strogatz, 1998), in which a highly connected ‘rich club’ balances the metabolic cost of maintaining long-range projections with neural efficiency (Bullmore and Sporns, 2012). These ‘hub’ neurons with long range projections will likely need to have more activity. Unlike the projection neurons from the perirhinal cortex to the medial prefrontal cortex, however, the neurons that formed the reciprocal projection to perirhinal cortex were not more likely show age-related differences in behaviorally induced activation. This observation points to an overall vulnerability of the prelimbic cortex in advanced age that is not influenced by efferent connectivity to the perirhinal cortex. One possibility that remains to be examined is whether the prelimbic neurons that receive afferent input from the PER are more likely to show age-related alterations in activity dynamics. Future work will need to use approaches for transynaptic anterograde tracing (Zingg et al., 2017) to explore this possibility.

Importantly, in both the deep and superficial layers, the percent of prelimbic cortical projection neurons to the PER that expressed *Arc* correlated with WM/BAT performance. This observation supports a growing body of evidence that bidirectional functional connectivity between the prelimbic and perirhinal cortices is critical for higher cognitive function and vulnerable in advanced age. It also indicates that the increased probability of *Arc* expression in projection neurons is not likely an artifact of the presence of the retrograde tracer CTB altering immediate-early gene transcription. Interestingly, in the superficial layers of prelimbic cortex a relationship between neuron activation and WM/BAT performance was only evident within the population of cells that projected to PER and not when cells without CTB labeling were included in the analysis. This finding suggests that the prelimbic efferent projections to the PER are particularly important for WM/BAT performance.

When the population overlap of active neurons across the WM/BAT and continuous spatial alternation tasks was evaluated, we found that the probability a cell would express *Arc* during both behavioral epochs did not differ between young and aged rats. These findings differ from our previous results, which showed a lower similarity score for prelimbic cortical neurons (that is, less population overlap) in aged animals compared to young (Hernandez et al., 2018b). These differences can likely be attributed to differences in behavioral performance between the two studies, particularly for the aged rats that made significant more errors in the current study relative to Hernandez et al. (2018b). Notably, the similarity scores reported here for both age groups are similar to those reported for the young rats in Hernandez et al. (2018b), indicating the recruitment of two distinct neuronal ensembles that distinguish between the different behavioral epochs.

Together, these data indicate that there are unique patterns of prelimbic cortical neuron activity in young and aged rats that vary across behavioral states. While the aged rats show greater activation in the home cage, they are less able to recruit prelimbic cortical networks during behavior. Importantly, the aged rats that do show greater activation of prelimbic cortical neurons during behavior also performed better on the WM/BAT task. Thus, maintaining prelimbic cortical activity dynamics and functional connectivity between this structure and the perirhinal cortex may be critical for sustaining cognition in advanced age.

## Acknowledgements

This work was supported by the McKnight Brain Research Foundation, National Institute on Aging (R01AG049722 and R21AG051004 to SNB, and F31AG058455 to ARH), and University of Florida, College of Medicine Scholars Program Awards (LMT and MEB).

## Notes

### Competing Interest Statement

The authors have declared no competing interest.

